# Estimating flux dynamics from metabolite concentration time courses with dynafluxr

**DOI:** 10.1101/2025.07.22.666125

**Authors:** Serguei Sokol, Svetlana Dubiley, Pauline Rouan, Cyril Charlier, Guy Lippens, Pierre Millard

## Abstract

**Motivation:** Quantifying metabolic fluxes and their dynamics is crucial for addressing fundamental and applied questions in systems and synthetic biology, health, and biotechnology. Since metabolic fluxes are not directly observable, they must be estimated using model-based approaches. While several methods are available to quantify steady-state fluxes, approaches for quantifying flux dynamics remain limited.

**Results:** We present a method to quantify flux dynamics from metabolite concentration time courses. Our approach employs constrained B-splines to solve mass balance equations without requiring prior knowledge of catalytic or regulatory mechanisms, or enzymatic parameters. We provide a comprehensive mathematical framework for dynamic flux calculation and demonstrate that our method enables accurate and robust quantification of flux dynamics for both simulated and experimental datasets.

**Availability and implementation:** `dynafluxr` is implemented in R and available on Windows, Unix, and MacOS platforms. The source code and the documentation are freely distributed under GPL-3 license at https://github.com/MetaSys-LISBP/dynafluxr/. A companion notebook producing figures similar to those of this article is available at https://github.com/MetaSys-LISBP/DynafluxR_notebook.

## Introduction

Quantifying metabolic fluxes is central to understand the function and regulation of living organisms, as well as to engineer efficient biological systems [11, 15, 3, 18, 26]. Steady-state (i.e. static) fluxes can be quantified by flux balance analysis (FBA) which relies on stoichiometric models and inequality constraints [7]. Static fluxes can also be calculated by ^13^C-metabolic flux analysis (^13^C-MFA) where underdetermination is compensated by isotope balance equations to close the stoichiometric system [1]. The practical value of these approaches is recognized and they are widely used in systems and synthetic biology.

However, many questions require metabolic fluxes to be monitored dynamically. This can be achieved through kinetic models of metabolism, which require, in addition to the network topology, extensive knowledge of enzyme function including detailed information on catalytic and regulatory mechanisms to determine the most appropriate rate laws. These models also depend on enzymatic parameters (such as *K*_*m*_ and *K*_*I*_) and thermodynamic parameters (*K*_*eq*_), which can be measured or estimated from experimental data. However, building accurate kinetic models of the dynamics of metabolic systems [14, 28] is data- and resource-intensive, and difficult to scale for large networks because of computational and experimental bottlenecks [25].

While an intermediate approach based on splines or Kalman filtering has been proposed to estimate the dynamics of growth rates and extracellular uptake or production fluxes [5], this approach cannot provide information on the dynamics of internal fluxes. This particular issue is partially resolved in dynamic MFA by extending the approach used to calculate the metabolite dynamics to internal fluxes, which are modeled using B-spline functions and nonlinear optimization [2, 13, 27]. However, this comes at the price of assuming stationary state to internal metabolite levels, with only external metabolite levels left unbalanced and free to vary over time. An approach for estimating complete flux dynamics including unbalanced internal metabolites was introduced in [8], using stoichiometric modeling and slope estimation at each time point to describe the point-wise time evolution of fluxes. The limitations of this proof-of-principle were however that no detailed mathematical framework was provided, missing data typically encountered in real-world experiments were not addressed, uncertainty propagation was overlooked, and no software implementation was proposed.

Here, we present a detailed mathematical framework to estimate flux dynamics from metabolite concentrations time-course, based only the matabolic network topology. We first describe the mathematical background of the proposed algorithm to calculate fluxes and estimate uncertainties propagation. We also highlight key considerations for implementation, which are incorporated into the accompanying software, `dynafluxr`. We demonstrate the applicability and robustness of our approach using a synthetic dataset and present a real-word use-case in which fluxes are estimated in a multi-enzymatic pathway monitored by real-time NMR spectroscopy.

## Algorithm

### General principles

The main steps of the algorithm are outlined in Figure 1. The proposed approach is based on the use of constrained B-splines to fit metabolite time courses with stoichiometric constraints. As detailed in the next section, the advantages of constrained B-splines are that they respect the positivity and, if necessary, monotonicity of metabolite time courses, properties that are difficult to achieve with other basis functions such as Fourier series or wavelets. The coefficients of the fitted B-splines (which we call measured coefficients) are fitted in a linear system based on the stoichiometric matrix. The principal unknowns in this linear system are the B-spline coefficients of the reaction rates which are estimated directly by constrained least squares optimization, with inequality constraints on the positivity and monotonicity of the simulated metabolite time curses and equality constraints, e.g. in the form of zero starting values for absent metabolites at the start of experiments.

**Fig. 1.**
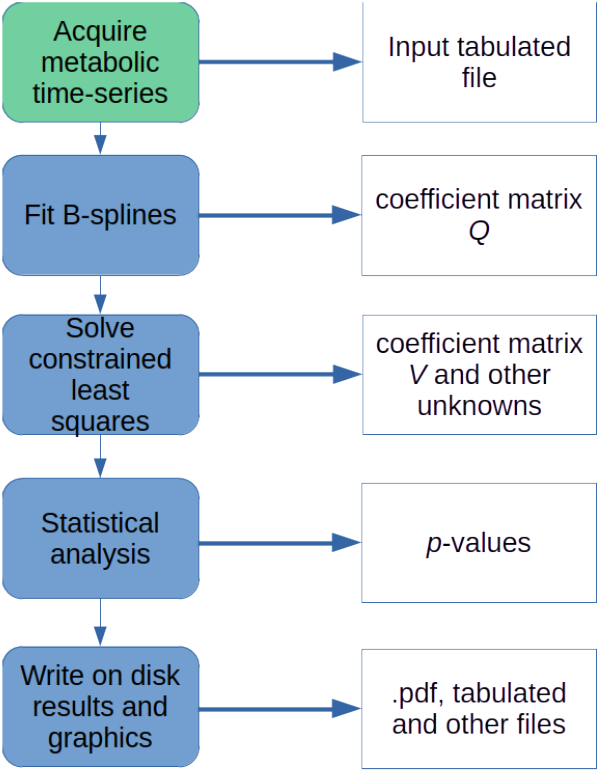
Workflow of the `dynafluxr` package. The green box is the experimental step and the blue boxes represent software tasks. The order of the tasks in the workflow is indicated by thin vertical arrows. The thick horizontal arrows point to the main results of each step (white boxes). Matrices are described in the text.

The following sections describe the advantages and drawbacks of two possible approaches to modeling time curses : the integrative approach, where the metabolite dynamics coefficients are fitted directly and the differential approach, where the fit is performed on the coefficients of the first derivatives of the metabolite time courses.

### Describing metabolite time courses with constrained B-splines

Let us first recall the fundamentals of B-splines. B-splines are piecewise polynomial functions [6] that are widely used for interpolation in industry and research. B-spline basis functions *B*_*k,n*_ of degree *n* typically ressemble a series of bell-curves (Figure 2) and can be recursively defined over a sequence of knots {*τ*_*k*_, *k* = 1, …, *K*, s.t. *τ*_*k*+1_ ≥ *τ*_*k*_, ∀*k*}:

**Fig. 2.**
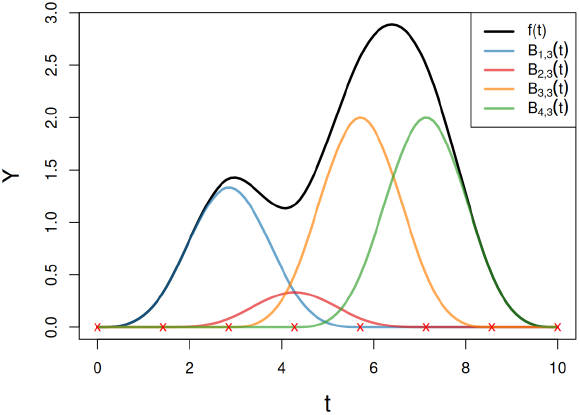
Example of B-spline approximation. The black line is a function *ϕ*(τ), represented as a sum of weighted B-splines (colored lines) defined over knotsrepresented here by red crosses.

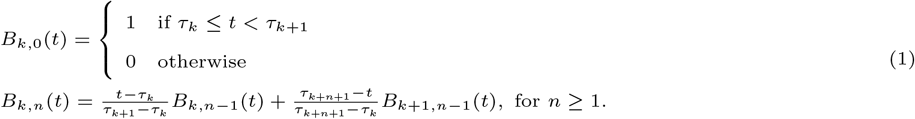

Any real-valued time-varying function can be approximated as a weighted sum of basis functions *f* (*t*) = ∑_*k*_ *q*_*k*_*B*_*k,n*_(*t*) where {*q*_*k*_, *k* = 1, …, *K* − *n* − 1} are B-spline coefficients or weights.

It follows from the definition of B-splines (Equation 1) that *B*_*k,n*_(*t*) *≥* 0, ∀*k, n, t*. If moreover, all *q*_*k*_ *≥* 0 then *f* (*t*) *≥* 0, ∀*t*. We have used this property in the R package bspline [23] to achieve constrained B-spline fitting with non-negative *q*_*k*_ coefficients.

This key feature ensures that the fitted chemical species concentrations are always non-negative.

Derivative and integration operators can also be defined based only on B-spline coefficients. If *f* ^*′*^(*t*) = ∑_*k*_ *p*_*k*_*B*_*k,n* − 1_(*t*), the first derivative of *f* (*t*), then the coefficients *p*_*k*_ and *q*_*k*_ are related by

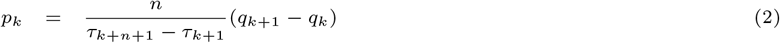

or in matrix form

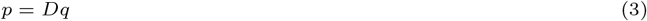

where *D* is an appropriately sized differentiation matrix whose coefficients are given by Equation 2.

Assuming that the coefficients *p*_*k*_ of *f* ^*′*^(*t*) are known, B-spline curves can be integrated to retrieve the coefficients *q*_*k*_ of *f* (*t*):

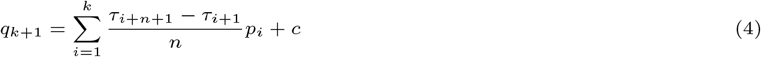

or in matrix form

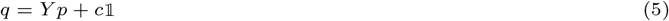

where *Y* is an integration matrix of appropriate size whose coefficients are defined by Equation 4, *c* is an arbitrary context-dependent constant, e.g. the initial condition, *f* (0), and 𝟙 is a vector of ones.

Another useful feature of the bspline package is that the fitted curves can be constrained to be monotonous, i.e. *q*_*k*+1_ *≥ q*_*k*_, ∀*k*. It then follows from (Equation 2) that in this case *p*_*k*_ *≥* 0, ∀*k* which means that *f* ^*′*^(*t*) *≥* 0, ∀*t*, hence *f* (*t*) is monotonically increasing. The same property can be exploited for monotonically decreasing functions. These constraints can be useful for species that are known to be only consumed (e.g. monotonically decreasing substrate concentrations) or only produced (e.g. monotonically increasing concentrations of end-products).

### Differential least-squares formulation

The bspline packagecan be used to fit individual metabolite time course *M*_*j*_ (*t*) (the first step in Figure 1). This preliminary fit does not take any system level information into account since the stoichiometric matrix is not used at this stage. The goal is to capture the dynamics of the time courseswith the coefficients of their first derivative subsequently fitted by least-squares optimization involving the stoichiometric matrix. The fitted B-spline functions are constrained to be non-negative and, if appropriate, monotonic.

The continuous time evolution of a given metabolite, *M*_*j*_ (*t*), can be expressed in terms of fourth order B-splines as follows:

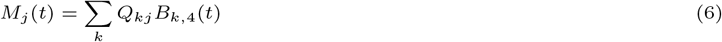

where *Q* is the coefficient matrix from the preliminary fitting step. Forth order B-splines are used to ensure that reaction rates be modeled with third order B-splines, a high enough order in most cases to model complex dynamics with smooth curves and continuous second derivatives.

The *j*-th column *Q*_.*j*_ corresponds to metabolite *M*_*j*_ (*t*), thus the *k*-th element *Q*_*kj*_ corresponds to *k*-th coefficient of the basis function *B*_*k*,4_(*t*). The knot sequence (the same for all metabolites) is a regularly spaced time grid covering the entire measurement interval and determined by the number of internal knots (user-defined, 5 by default). This number should be set to the lowest value at which satisfactory fits are obtained. In our applications with 5 to 1000 time points, accurate fits were obtained with 0 to 5 internal knots. Using too many internal knots can lead to over-fitting and non-physical results such as excessive oscillations. Note that the fitting process is robust to the presence of a few missing data, provided that the number of available data points is higher than the number of B-spline coefficients and that each knot interval contains at least one valid data point. This allows the analysis on incomplete datasets where the dynamics of the system have only partially been measured.

Note that metabolites that are not detected experimentally, for which all data are missing, are simply excluded from the least-squares cost functions. This point is considered in greater detail later in the paper.

B-splines have already been used to fit metabolite dynamics [8, 13] but the originality of our approach, described in the following paragraphs, is to perform a constrained least-squares fitting problem directly on B-spline coefficients of reaction rates *V*.

Let *S*_*jl*_ be an element of the stoichiometric matrix, *S*, corresponding to metabolite *M*_*j*_, which is either produced or consumed (depending on the sign of the product *S*_*jl*_*v*_*l*_) by an *l*-th reaction at a rate *v*_*l*_. We formally distinguish the reaction rate, *v*_*l*_, from the metabolic flux, *f*_*jl*_ = *S*_*jl*_*v*_*l*_, induced by this reaction. In this context, reaction rates indicate how often the corresponding reactions take place per unit time (dimension of 1*/*[Time]). In contrast, the metabolic flux, *f*_*jl*_, represents th amount of *M*_*j*_ transformed by the *l*-th reaction per unit time (dimensions of [Concentration]*/*[Time]). When |*S*_*jl*_| = 1, *v*_*l*_ and *f*_*jl*_ have the same absolute values (which explains why they are often considered synonyms) but the two quantities vahe different physical meanings. The overall metabolic flux for *M*_*j*_ can then be calculated from 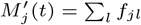, or over the whole concentration vector *M* (*t*), using

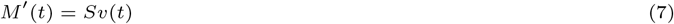

where *v* is a vector of unknown reaction rates. This well-known stoichiometric ordinary differential equation represents the mass conservation law at every time point *t*.

Modeling *v*_*l*_(*t*) as ∑_*k*_ *V*_*kl*_*B*_*k*,3_(*t*) where *V*_*kl*_ are third order B-spline coefficients, the first derivative of *M*_*j*_ can be written

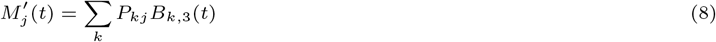

and the *P*_*kj*_ coefficients can be obtained from *Q*_*kj*_ based on (Equation 3)

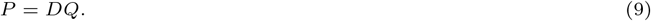

Substituting these two expressions for 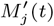 and *v*_*l*_(*t*) into Equation 7 and simplifying *B*_*k*,3_(*t*)yields the following matrix equation for the B-spline coefficients *V* :

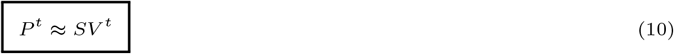

where the superscript ^*t*^ represents the transposition operator. In this form, the system can be solved by classical linear algebra methods based on generalized inverses.

The Equation 10 has a unique solution if and only if (i) *S* is a “tall” matrix, i.e. the number of rows, *n*_*r*_, (the number of species in the vector *M* (*t*)) is equal to or greater than the number of columns, *n*_*c*_ (the number of reactions in *v*(*t*)), and (ii) *S* is of full rank, i.e. rank(*S*) = *n*_*c*_. The condition *n*_*r*_ ≥ *n*_*c*_ must still be satisfied after the rows of *S* and columns of *Q* and *P* corresponding to unmeasured metabolites are removed. Otherwise, the system is under-determined and has an infinite number of solutions. In this situation, the software provides the solution of minimum norm [9], as used for example in parsimonious FBA [20] with rational that since high fluxes correspond to higher enzyme turnover, and since producing enzymes is costly, cells have evolved to minimize overall flux.

Another situation that can arise in practice is for one or several metabolites to only have relative concentrations available, as typically occurs when metabolites are analyzed by mass spectrometry without an internal standard, or by nuclear magnetic resonance when the number of atoms contributing to the signal cannot be determined. In this case, additional scaling factors {*α*_*j*_}

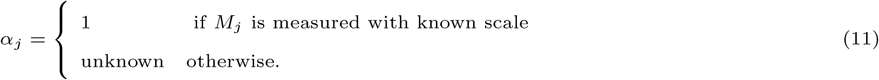

The corresponding metabolite measurements are then scaled as

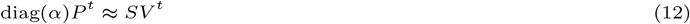

where diag(*α*) is a diagonal matrix with vector *α* on the main diagonal. Since this scaled system is difficult to represent in the classical form, *Ax* = *b*, with an explicit matrix *A*, right hand side *b* and unknown vector *x* enclosing *α* and *V*, it is solved using the generalized minimum residual (GMRES) method [19, 24], which does not require an explicit matrix formulation.

Differential least-squares (DLS) optimization is simple, elegant and computationally efficient, but cannot include constraints on initial values, for example the values measured for a set of metabolites at the beginning of the experiment which are crucial for successful fitting in some cases. This is the main motivation for developing the integral least-squares (ILS) method described in the following section.

### Integral least-squares formulation

*Sv*(*t*) can be integrated over time using Equation 5 to obtain a linear system in the least-squares sense:

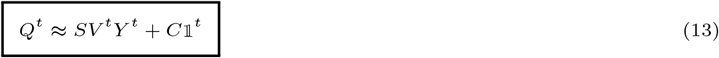

where *C* is an appropriate vector of additive constants, one per metabolite. The unknowns in this equation are the matrix *V* and the vector *C*. Inequality constraints for possible monotonicity

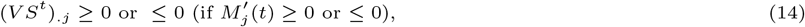

for non-negativity

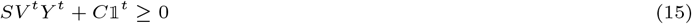

and equality constraints on initial values (too cumbersome to be detailed here) can be added to obtain the ILS system. In this case, the solution will require more advanced methods [12, 22] than classical inverses. Unknown scaling factors *α* can also be included, as in the DLS system. ILS is the default optimization method in `dynafluxr`, despite being more computationally intensive, because it allows the inclusion of initial values and other types of constraints.

Optimization by ILS or DLS corresponds to the second blue box in Figure 1.

### Sensitivity analysis

The package `bspline` provides covariance matrix estimates of the uncertainties of the fitted B-spline coefficients for each metabolite. The covariance matrix for the *j*-th metabolite is represented as the product of a basis covariance matrix, *Z*^*q*^, and a corresponding factor 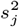:

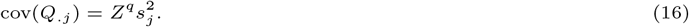

The same basis matrix *Z*^*q*^ is used for all fitted metabolites. The vector of 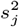 elements is referred as *s*^2^ hereafter. Note that elements on the same row in *Q* are not correlated as they correspond to independently measured metabolites.

To calculate the covariance matrix for *V*, solution of the DLS optimization problem, we first calculate the covariance of *P*_.*j*_ :

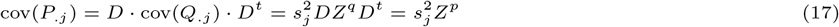

where *D* is differentiation matrix introduced in equation 3 and *Z*^*p*^ is a shorthand expression for *DZ*^*q*^*D*^*t*^.

The covariance of any two elements of *V* can then be calculated using

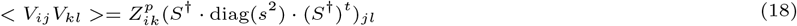

where diag(*s*^2^) is a diagonal matrix with the elements of vector *s*^2^ on the main diagonal. The derivation of this formula is given in SupplementaryFile1.

The square root of *< V*_*kl*_*V*_*kl*_ *>* is the standard deviation (SD) of the *k*-th B-spline coefficient for the *l*-th reaction rate. This formula can be used to trace B-spline curves with smooth confidence intervals.

For ILS, the formula for estimating uncertainties is too complex to be presented here but the resulting method is implemented in `dynafluxr` package (see source code for details).

### Goodness-of-fit evaluation

Goodness of fits in `dynafluxr` are estimated using *χ*^2^-tests. The statistic for the *j*-th species :

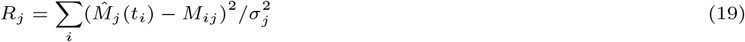

is the sum of squared residuals divided by the variance 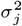 estimated from the variance of differences between fitted and measured values: 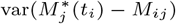. Note that *M*_*ij*_ are measurements and *M* ^*∗*^(*t*) are preliminary fitted B-splines (the superscript ^*∗*^ means that corresponding fit was performed on a grid with an optimal number of knots estimated automatically in `dynafluxr`), while 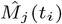 in Equation 19 is the value of splines obtained (i.e. estimated) by integrating *Sv*(*t*). If the uncertainties in *M*_*ij*_ are normally distributed (a common assumption) then *R*_*j*_ follows a *χ*^2^ distribution with *n*_*m*_ − *n*_*k*_ degrees of freedom (where *n*_*m*_ is the number of measurements and *n*_*k*_ is the number of B-spline coefficients estimated for a given metabolite). The test’s *p*-value is then *P* [*χ*^2^(*n*_*m*_ − *n*_*k*_) ≥ *R*_*j*_]. Since the null-hypothesis of this test is that *R*_*j*_ follows *χ*^2^(*n*_*m*_ − *n*_*k*_), small *p*-values (typically defined as *p* ≤ 0.05) indicate poor quality fit.

Note that these *χ*^2^ tests are performed globally but also separately for each species, allowing user to differentiate between well-fitted and poorly fitted metabolites.

Another point to keep in mind is that *R*_*j*_ can be nonzero even in the absence of uncertainty, e.g. for synthetic data. This is because polynomials such as B-splines cannot perfectly fit other types of functions such as exponential and sigmoid functions.

Goodness-of-fit testing corresponds to the third blue box in Figure 1.

## Implementation

The algorithms described in the previous sections are implemented in the R [17] packages `bspline` and `dynafluxr`, accessible from CRAN [23, 21]. The `dynafluxr` package can be used as a command line tool, programmatically, as a classical R package or as a local Shiny [4] application with a graphical user interface on a dynamic web page. The input data (network topology and measured metabolite concentrations) can be provided as tab-separated values (TSVs). The results are written in TSV files and PDF plots are generated for visual inspection. Results can also be returned as R objects for further manipulation in R (see the example `notebook.Rmd` at https://github.com/MetaSys-LISBP/DynafluxR notebook). In this article, we have used `dynafluxr` v1.0.1 and `bspline` v2.5.0.

As for other CRAN packages, a user manual and step-by-step description of a use-case (human readable text explaining R code excerpts with embedded results) are provided. The source code of these packages is available under open source GPLv3 licence from the following GitHub repositories https://github.com/MathsCell/bspline and https://github.com/MetaSys-LISBP/dynafluxr.

## Validation using a synthetic dataset

The ability of the proposed method to accurately retrieve reaction rates was first tested using simulated data. We built a kinetic model of the network shown in Figure 3A, consisting of eleven metabolites (A to K) and nine reactions (r1 to r9). Reaction r1 was regulated by the law of mass action, while the other reactions were modeled using Michaelis-Menten kinetics, with reaction r2 regulated by metabolite G through competitive inhibition. Reaction rates and metabolite dynamics were simulated using the software COPASI [10] at 20 time points equally spaced over 25 min. This model has been uploaded to the Biomodels database (under identifier MODEL2502210001). The stoichiometric model as well as the simulated metabolite time courses can be found in Supplemental Data as files `network_toy.txt` and `data_toy.tsv` respectively. The resulting fluxes and metabolite concentrations show a broad range of dynamic profiles. For instance, some metabolites are continuously depleted (e.g., A), some only accumulate (e.g., H and E), and others increase in concentration at first before decreasing (e.g., G and F).

**Fig. 3.**
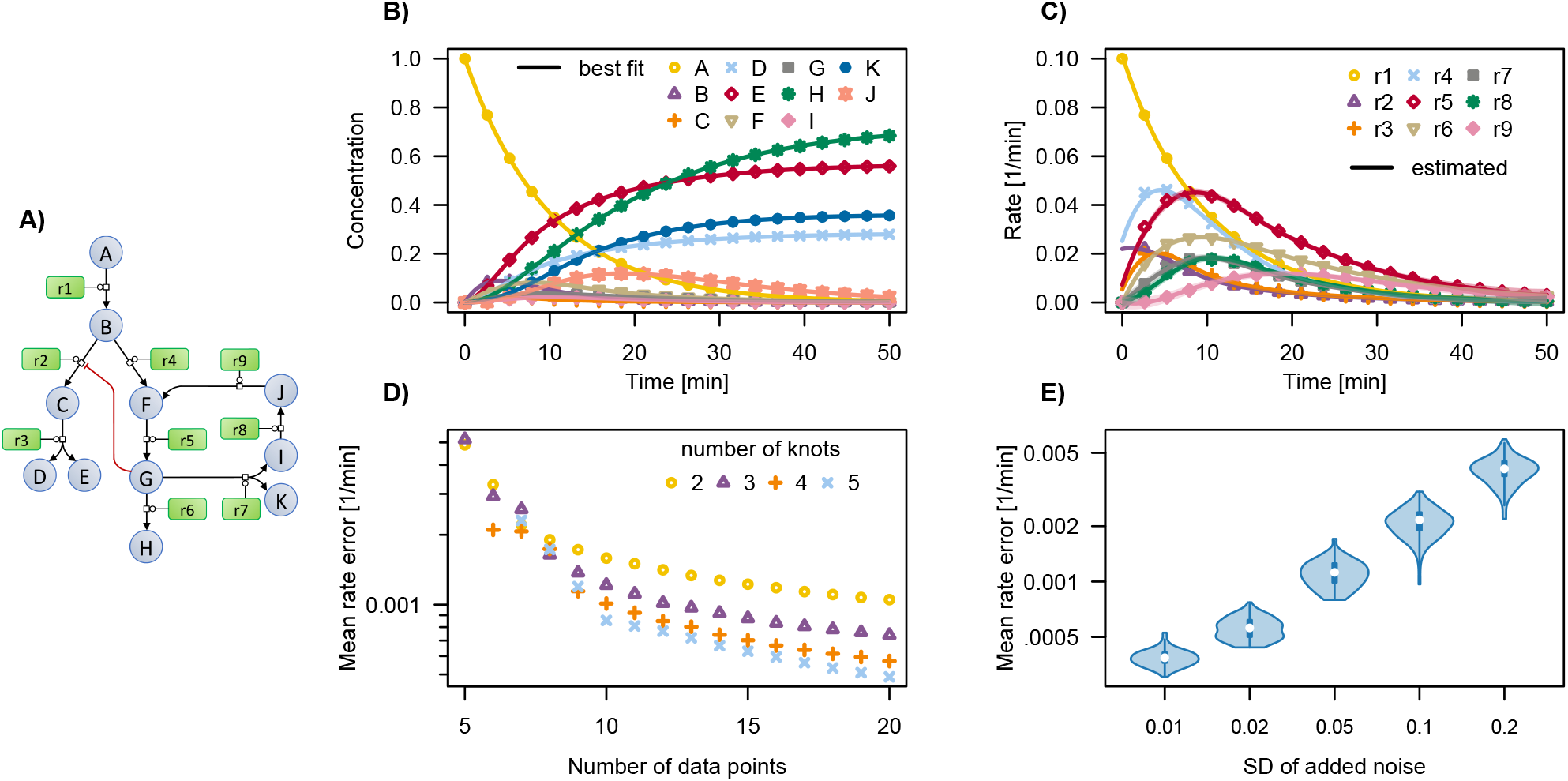
Validation of `dynafluxr` with simulated data. (A) Topology of the metabolic network used for simulations. (B,C) Simulated and estimated (B) metabolite and (C) flux dynamics with simulated data as points and `dynafluxr` best fits shown as lines. (D,E) Mean error on reaction rates (D) as a function of the number of time points and knots used for flux calculation and (E) as a function of the added noise level.

We used `dynafluxr` to estimate flux dynamics from the simulated metabolite dynamics, using only the network stoichiometry and default parameters. The calculations took less than 1 s to perform on a laptop computer. All metabolite dynamics were accurately captured, with a mean error in metabolite concentrations of 2.9 · 10^−4^ (Figure 3B). All reaction rates were also estimated with high accuracy (Figure 3C), with a mean error of just 4.9 · 10^−4^, very low relative to the flux values. These results confirm that under ideal conditions (when measurements are noise-free, all metabolites are measurable, and the network topology is correctly defined) the proposed method does indeed provide accurate flux estimates.

To assess the robustness of our method with respect to the number of measured time points and the number of knots, we calculated fluxes from the same simulated dataset with 5 to 20 equally spaced time points, using 2 to 5 knots. Figure 3D shows that the mean reaction rate error was high when less than 10 time points were included. In situation such as this, estimates with fewer knots are more accurate because of instabilities when the number of knots is similar to the number of measurements. When at least 10 time points were included however, the error stabilized at *≤* 10^−3^ regardless of the number of knots used. In these cases, using more knots slightly improves the accuracy of the predicted fluxes, providing more flexibility to accurately fit the metabolite dynamics. These results indicate that above a certain data threshold (at least 7-10 time points in this example), our approach is robust with respect to the number of data points and knots used. Note that since the optimal number of knots and measurements depend both on the metabolic network and on the metabolite dynamics, optimal fitting parameters should be determined on a case-by-case basis.

Flux calculations were also repeated with increasing levels of Gaussian noise, ranging from 1 % to 20 % of the mean concentration of each metabolite. All datasets were successfully fitted, and, as expected, the mean error increased gradually with the noise level (Figure 3E). In this example, the errors on reaction rates remained low for noise SDs below 10 %. Although the precise effects of noise depend on the studied metabolic network and metabolite dynamics, these results confirm that our method is robust to noisy datasets.

## Application to a multi-enzymatic system

We further evaluated `dynafluxr` by calculating fluxes in a multi-enzymatic system reproducing the first three reactions of glycolysis (Figure 4A). As described in [16], the experiment involved adding glucose (5 mM) at *t* = 0 to a buffer solution containing ATP (10 mM) and the first three glycolytic enzymes: glucokinase (HXK), glucose-6-phosphate isomerase (PGI), and phosphofructokinase (PFK). The concentration time courses of glucose (GLC), glucose-6-phosphate (G6P), fructose-6-phosphate (F6P), and fructose-1,6-bisphosphate (FBP) were monitored by real-time ^1^H-NMR. Data were recorded as a pseudo-2D spectrum with 1,024 equally spaced time points from 10 to 723.8 min after the glucose pulse. The integrals of the signals were obtained with MultiNMRFit [16]. Absolute concentrations of GLC and G6P were determined by normalization against the internal standard. For F6P and FBP, only relative concentrations could be determined because of assignment ambiguity. The stoichiometric model as well as the measured data can be found in Supplemental Data as files `network_glyco.txt` and `data_glyco.tsv` respectively.

**Fig. 4.**
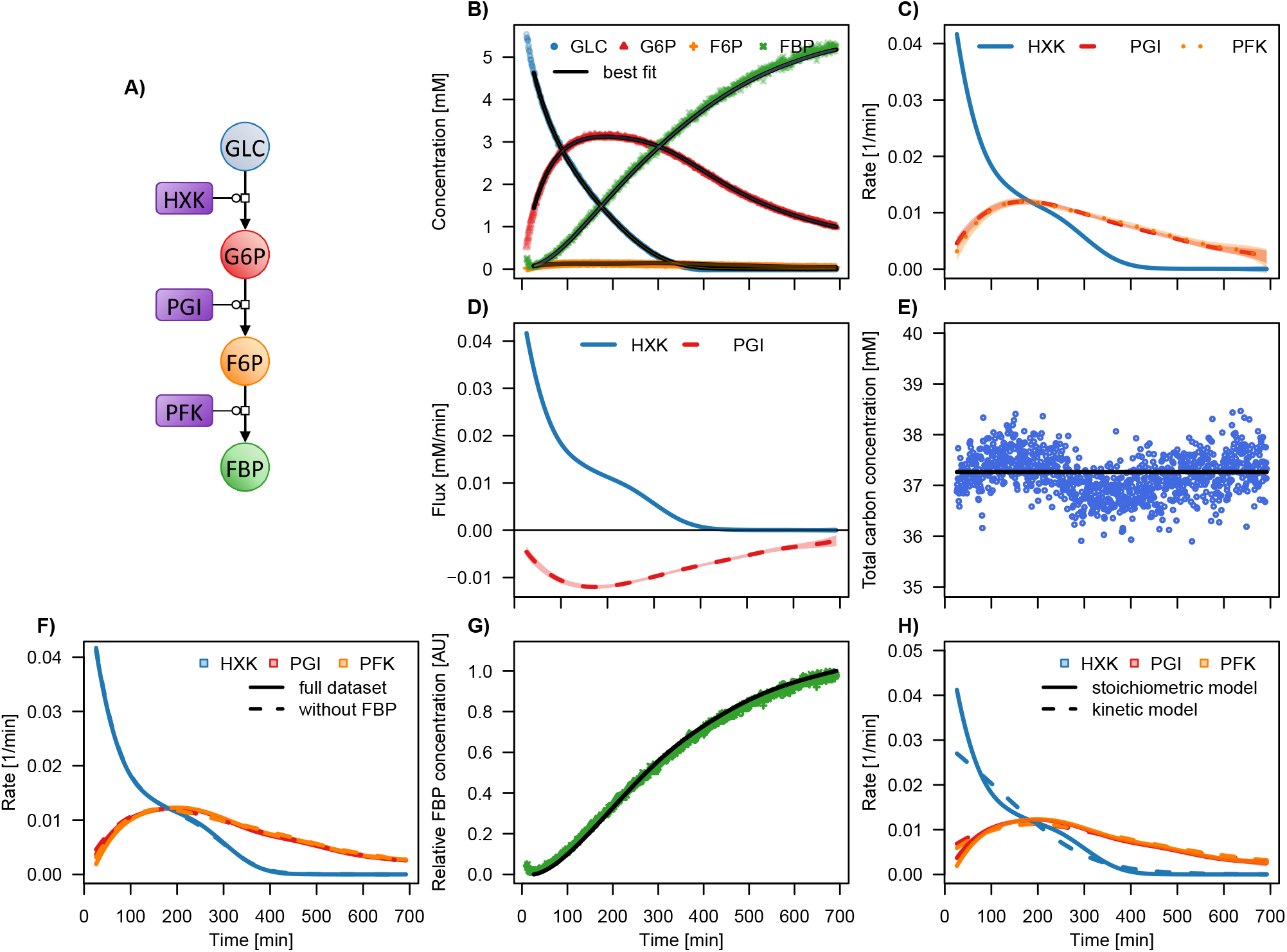
Flux analysis in a multi-enzymatic reaction system. (A) Topology of the metabolic network used for simulations. (B) Measured and estimated metabolite dynamics, with experimental data shown as points and `dynafluxr` best fits as lines. (C) Reaction rates calculated with `dynafluxr`. (D) Contribution of HXK and PGI fluxes to G6P dynamics. (E) Experimentally measured (circles) and estimated (line) carbon balance. (F) Comparison of reaction rates estimated from the complete metabolite dataset (solid lines) and after excluding FBP (dotted lines). (G) The predicted FBP concentration, whose evolution was not used for flux calculations, is shown as a black line and experimental data are shown as dots. (H) Comparison of reaction rates estimated by `dynafluxr` and by COPASI (using a kinetic model). The colored bands around the fitted curves in parts (B), (C) and (D) correspond to *±*2 standard deviations (they are in some cases indistinguishable from the curves themselves).

Experimental metabolite time courses are shown in Figure 4B. For GLC and G6P, absolute concentrations were fitted directly, while for F6P and FBP, scaling factors were used because only relative concentrations were available. Reaction rates (Figure 4C) were estimated in less than 1 s, and the complete dataset was satisfactorily fitted with 95 % confidence. The HXK flux was maximal at the start of the experiment, when the GLC concentration was highest, and decreased over time as substrate levels declined. In contrast, fluxes through PGI and PFK initially increased as G6P and F6P concentrations rose, before decreasing as GLC was completely converted into FBP. The estimated rates allowed us to isolate the time-resolved contributions of individual reactions to metabolite dynamics (see Figure 4D for the contribution of HXK and PGI to G6P kinetics), providing detailed insights into system dynamics. `Dynafluxr` also validated the self-consistency of the dataset by confirming the carbon balance (Figure 4E).

The number of available measurements for the considered network meant that the system of equations was overdetermined. To test the robustness of our approach to missing data, we removed FBP measurements from the dataset and used `dynafluxr` to (i) calculate fluxes using the reduced dataset and (ii) predict the concentration time course of FBP. The flux profiles with reduced dataset remained similar to those obtained with the complete dataset (Figure 4F), and the predicted FBP dynamics closely matched experimental data (Figure 4G). These results highlight the robustness of `dynafluxr` in handling missing data and illustrate how metabolite profiles can be predicted. This feature may be useful to deduce spectral assignments by comparing predictions to the profiles of putative signals that cannot be assigned experimentally.

Finally, we compared our method with the current state-of-the-art in kinetic modeling approaches. A kinetic model of the three reactions was constructed, assuming Michaelis-Menten kinetics, with HXK and PFK as irreversible reactions and PGI as reversible. The model has been deposited in the Biomodels database (under identifier MODEL2502210002). The 18 parameters (10 kinetic constants, the initial concentrations of six metabolites, and two scaling factors for F6P and FBP) were estimated using COPASI by fitting the complete dataset with particle swarm optimization (5,000 iterations, swarm size of 50). The calculations were completed within a few minutes. The fluxes simulated using the best-fitted COPASI parameters were similar to those obtained using `dynafluxr` (Figure 4H). While the results of this test are not generalizable to all experimental scenarios, they nevertheless highlight the efficiency and simplicity of `dynafluxr`, which is less demanding in computational resources and assumptions than kinetic modeling. `Dynafluxr` also provides confidence intervals of the predicted fluxes, which are difficult to calculate for kinetic models.

## Conclusions

Current flux calculation approaches are based either on stoichiometric models, which are easy to construct and computationally cheap to analyze but inherently static, or on kinetic models, which are dynamic by definition but challenging to construct and parameterize. We have developed a method that combines the strengths of these two approaches to estimate flux dynamics from metabolite concentration time courses.

This method is intrinsically dynamic, but relies on minimal biological assumptions (only the network topology) and does not require information on rate laws, catalytic or regulatory mechanisms, or kinetic parameters. Moreover, our method combines a comprehensive mathematical framework with many practical features: computational efficiency resulting from the linear formulation of the problem; amenability to uncertain or missing data; enforced non-negativity and monotonicity of metabolite dynamics; and integration of absolute and relative concentrations through the use of scaling factors.

The precision of the estimated fluxes depends on the condition of the stoichiometric matrix *S* and on the accuracy, precision and frequency of the metabolite concentration measurements. The results presented here for simulated data show the method is accurate and robust against measurement noise. It was also successfully applied to real-world data from real-time NMR experiments on a multi-enzymatic system. Satisfactory fits were obtained for both datasets, highlighting the simplicity and efficiency of the proposed approach.

The accompanying open-source software, `dynafluxr`, enables fast and reliable estimation of flux dynamics. Beyond addressing fundamental questions in metabolic systems biology, this information is useful in healthcare (e.g., to investigate metabolic diseases) and in biotechnology (e.g., to design more efficient metabolic systems). `Dynafluxr` can complement the current metabolic flux analysis toolbox and help build a reliable framework to better understand and control metabolic fluxes. It opens the way for real-time metabolic flux analysis and can support methodological developments, as illustrated here for data-driven metabolite annotation. Finally, since the proposed method is generic, it can directly be applied to other biological, chemical, geological, or physical systems.

## Author contributions statement

S.S. conceived the mathematical algorithm, programmed software, analyzed results, and wrote the manuscript;

P.M. conceived and supervised the whole project, analyzed NMR data and results, tested software, developed and programmed kinetic models, and wrote the manuscript;

S.D., P.R., C.C. and G.L conceived and conducted NMR experiments, analyzed the data, and reviewed the manuscript. PM and SS contributed equally to this work.

## Acknowledgments

We thank the MetaToul-Fluxomet platform for access to NMR facilities. This work was funded by the FUNCEMM (ANR-23-CE44-0038) and HEPATOTWIN (INRAE Metaprogram DIGIT-BIO) projects, and was also supported by MetaboHub (MetaboHUB-ANR-11-INBS-0010).

