## SupplementalData for "Estimating flux dynamics from metabolite concentration time courses with dynafluxr": Readme.md

This is supplementary data for a paper "Estimating flux dynamics from metabolite concentration time courses with dynafluxr" by Serguei Sokol, Svetlana Dubiley, Pauline Rouan, Cyril Charlier, Guy Lippens, and Pierre Millard.

BioRxiv preprint DOI : https://doi.org/10.1101/2025.07.22.666125

Files in this directory:

 * Readme.md: this file
 * SupplementaryFile1.pdf: formula derivation for uncertainty propagation;
 * network_toy.txt: stoichiometric model for simulated network;
 * data_toy.tsv: simulated data;
 * network_glyco.txt: stoichiometric model for upper glycolysis experiment;
 * data_glyco.tsv: measured data for upper glycolysis experiment.
