## SupplementalData for "Estimating flux dynamics from metabolite concentration time courses with dynafluxr": SupplementaryFile1.pdf

### Uncertainty formula derivation

Serguei Sokol

2025-06-24

This file is the SupplementaryFile1.pdf for a paper entitled "Estimating flux dynamics from metabolite concentration time courses with `dynafluxr`" by Serguei Sokol, Svetlana Dubiley, Pauline Rouan, Cyril Charlier, Guy Lippens, and Pierre Millard.

#### From metabolites to reaction rates

Using the same notations as in the paper, we recall that $Q$ is the matrix of B-spline coefficients for metabolites, covariance basis matrix $Z_q$ and $s^2$ vector of variance factors such that covariance matrix of column $j$ in $Q$ is given by

$$\mathrm{cov}(Q_{.j}) = s_j^2 Z^q$$

and covariance of $P_{.j}$ by

$$\mathrm{cov}(P_{.j}) = s_j^2 Z^p.$$

Now, we solve DLS problem to find $V$, matrix of rate coefficients:

$$V = P(S^\dagger)^t$$

where $S$ is stoichiometric matrix, $^\dagger$ means generalized inverse and $^t$ designates the transposition. For the sake of notation simplicity, we rewrite $S^\dagger$ as $A$. Note that the row components of $P$ are uncorrelated, like are the row components of $Q$, as they are corresponding to independent metabolite measurements.

We are looking for covariance of any two elements $V_{ij}$ and $V_{kl}$:

$$< V_{ij}V_{kl} >=< \sum_\alpha P_{i\alpha}A^t_{\alpha j}\sum_\beta P_{k\beta}A^t_{\beta l} >= \sum_{\alpha\beta} A_{j\alpha} < P_{i\alpha}P_{k\beta} > A^t_{\beta l} =$$

(as $P_{.\alpha}$ and $P_{.\beta}$ are decorrelated)

$$\sum_\alpha A_{j\alpha} < P_{i\alpha}P_{k\alpha} > A^t_{\alpha l} = \sum_\alpha A_{j\alpha}Z^p_{ik}s^2_\alpha A^t_{\alpha l} =$$

$$= Z^p_{ik}\sum_\alpha A_{j\alpha}s^2_\alpha A^t_{\alpha l} =$$

$$Z^p_{ik}(A\,\mathrm{diag}(s^2)A^t)_{jl}$$

Thus, covariance matrix of $V_{.j}$ can be calculated as

$$\mathrm{cov}(V_{.j}) = Z^p(S^\dagger \mathrm{diag}(s^2)(S^\dagger)^t)_{jj}.$$
